# Molecular Prognosis of Endometrial Adenocarcinoma by Expression Patterns of the V-ATPase C1 Subunit

**DOI:** 10.1101/2021.10.05.463029

**Authors:** Abdalla Dib Chacur, Juliana Couto-Viera, Anna Okorokova Façanha, Glenerson Baptista, Arnoldo Rocha Façanha

## Abstract

- Comparative analysis of expression patterns of *ATP6V1C1* encoding C1 subunit of V-H^+^-ATPase revealed that molecular alterations correlated with endometrial cancer of better prognosis were grouped with low *ATP6V1C1* expression while those correlated with worse prognosis were clustered with high *ATP6V1C1* expression levels.
- Expression patterns of C1 subunit in endometrial adenocarcinoma are associated with molecular malignancy signatures shared by histological subtypes of highest mortality rate suggesting that G3 adenocarcinoma exhibits molecular changes resembling endometrial serous carcinoma.
- *ATP6V1C1* might serve as novel prognostic marker allowing identification of targetable pathways for high-risk endometrial cancer.

## Main

Endometrial cancer (EC) was estimated to be the second most incident gynecological cancer in the world, and is classified into two subtypes, a low-grade Type I (UEC, uterine endometrioid carcinoma), the most common EC, usually estrogen dependent, carrying a good prognosis; and a Type II, including endometrial serous histological subtype (USC, uterine serous carcinoma), often not associated with increased estrogen exposure, carrying a poor prognosis [1]. Although usually diagnosed in early stages, the survival rate of UEC decreases when associated with worse prognosis criteria. Identification of molecular patterns related to an unfavorable evolution would allow a better understanding of the oncogenic process and, hence, individualized and more accurate therapies, preventing avoidable radical treatments involving aggressive surgery, radiation and multimodal chemotherapy.

The vacuolar/lysosomal H^+^-ATPase (V-ATPase), a multisubunit protein complex, is important for acidification-dependent degradation of tissue matrices allowing cell migration and plays a key role in pH regulation across epithelial cell layers, including during placentation, which involves intricate signaling, cell proliferation, and controlled invasion [2]. It was also shown that progesterone-induced decrease in uterine fluid pH involves the functional overexpression of the V-ATPase, as suggested by the expression patterns of subunits of the catalytic domain (A1 and B1/2), which were found to be highly expressed throughout the uterine luminal and glandular epithelia [3]. Epithelial expression of V-ATPase subunits in nonpregnant animals exhibits an apical abundance, however, as pregnancy proceed, the expression of these subunits became luminally pericellular, excepting in glandular epithelium, indicating a specific spatial-temporal membrane targeting redistribution of V-ATPases [2]. Furthermore, ectocervical cells acidify the vaginal canal by a mechanism of H^+^ pumping driven by V-ATPases located in cell apical plasma membranes, and this active net H^+^ secretion occurs constitutively throughout life, with acidification cycles up-regulated by estrogen [4]. Estrogen also induces proliferation of endometrial glands and tissues hyperplasia. In patients at risk, an accumulation of genetic damage led to proliferation of mutant clones under the mitogenic stimulus of estrogen, triggering histological changes that can culminate in malignant transformation [5].

Regarding these functional relationships of V-ATPases with the acid metabolism of female reproductive system, it is perhaps not surprising that specific modulations in functional expression of this pump has been associated with the acid metabolic reprogramming of different gynecologic cancers (e.g., [6]). However, the study of expression profiles of this enzyme is not trivial, since V-ATPases are composed of 13 subunits, divided into eight peripheral proteins grouped in a cytoplasmic V1 domain (A-H), responsible for the hydrolysis of ATP, and five intrinsic membrane proteins composing a V0 domain, represented by subunits a, c, c”, d and e. In mammals, some subunits of the enzyme have different isoforms. The C subunit which is important for the reversible structural and functional coupling of the V1 and V0 domains, has three isoforms in humans, namely C1, C2a and C2b [7].

The C1 isoform of subunit C of this oligomeric enzyme has been associated with an assembly of the holoenzyme of greater catalytic efficiency, related to a higher coupling between the V1 and V0 domains, and thus, the expression profile of this subunit has received increasing attention by its differential expression in different tumor cells (e.g., [8]), leading to a reprogramming of intra- and extracellular pH homeostasis [9,10], and even in nucleoplasm [11]. Disruption of cellular pH homeostasis is a hallmark of most types of cancer, regardless of their genetic or tissue origin [11]. A neutral-slightly alkaline intracellular pH is permissive to cell proliferation and evasion of apoptosis, whereas the decrease in extracellular pH favors the evasion of the immune response against the tumor, promotes the extracellular matrix remodeling and stimulates acid-activated proteases, facilitating the invasion and dissemination of tumor cells [12]. Tumor cell metastasis also depends on secreted lysosomal enzymes that participate in the extracellular matrix degradation and cell to cell detachment, which in turn, is related to altered expression profile of adhesion cell molecules. Lysosomal enzymes require a low pH for optimal activity, and in tumor cells V-ATPases regulate acidification of extracellular microenvironment [13].

Here, we used the Cancer Genome Atlas (TCGA) database to identify putative clustering expression patterns associated with the C1 isoform in UEC that could refine prognosis and identify targetable pathways for high-risk UEC. Genomic and clinical data on TCGA endometrial carcinoma were accessed through the CBioPortal (www.cbioportal.org)/ PanCancer database (n=529). The histological type selected was UEC (n=399), which represented 75.4% of the available cases, among these 397 had expression information by RNA-seq (RNAseq V2 RSEM), representing 99.5% of the samples. Levels of mRNA were evaluated for V-ATPase genes representing V1 domain and V0 domain, totalizing 25 genes. Clinical-pathological information was collected regarding mutation, staging, histological grade, percentage of invasion, survival; genetic alterations in V-ATPase genes expressed in UEC were analyzed and expression heatmaps of genes with similar patterns of *ATP6V1C1* expression were verified. Analysis of gene expression patterns according to histologic grade and significant genes coexpression was also performed. Comparison between two expression groups (*ATP6V1C1* of high >1 and low <-1 expression profiles) was performed in relation to the histological grade and the most incident genetic mutations. We also analyzed the correlation between the differential patterns of *ATP6V1C1* expression in relation to selected clustered expression genes and proteins. Statistical analysis was obtained from the cBioPortal platform, considered significant (p<0.005).

Data were analyzed according to the differential *ATP6V1C1* mRNA levels (high and low C1 isoform expression patterns). Remarkably, UEC grade 1 and 2 exhibited low expression of *ATP6V1C1*, while both G3 UEC and G3/high grade USC exhibited high *ATP6V1C1* expression (Suppl. Fig. S1 and Fig. 1a,d). Moreover, a few genetic alterations found in ATP6V1C1 correlated well with high C1 expression and G3 tumor grade (Suppl. Fig. S2). Analysis of only the G3 UEC showed better overall survival linked to medium ATP6V1C1 expression levels (Suppl. Fig. S2), suggesting that regular expression of V-ATPase isoform C1 is a significant factor influencing treatment responsiveness.

**Fig. 1.**
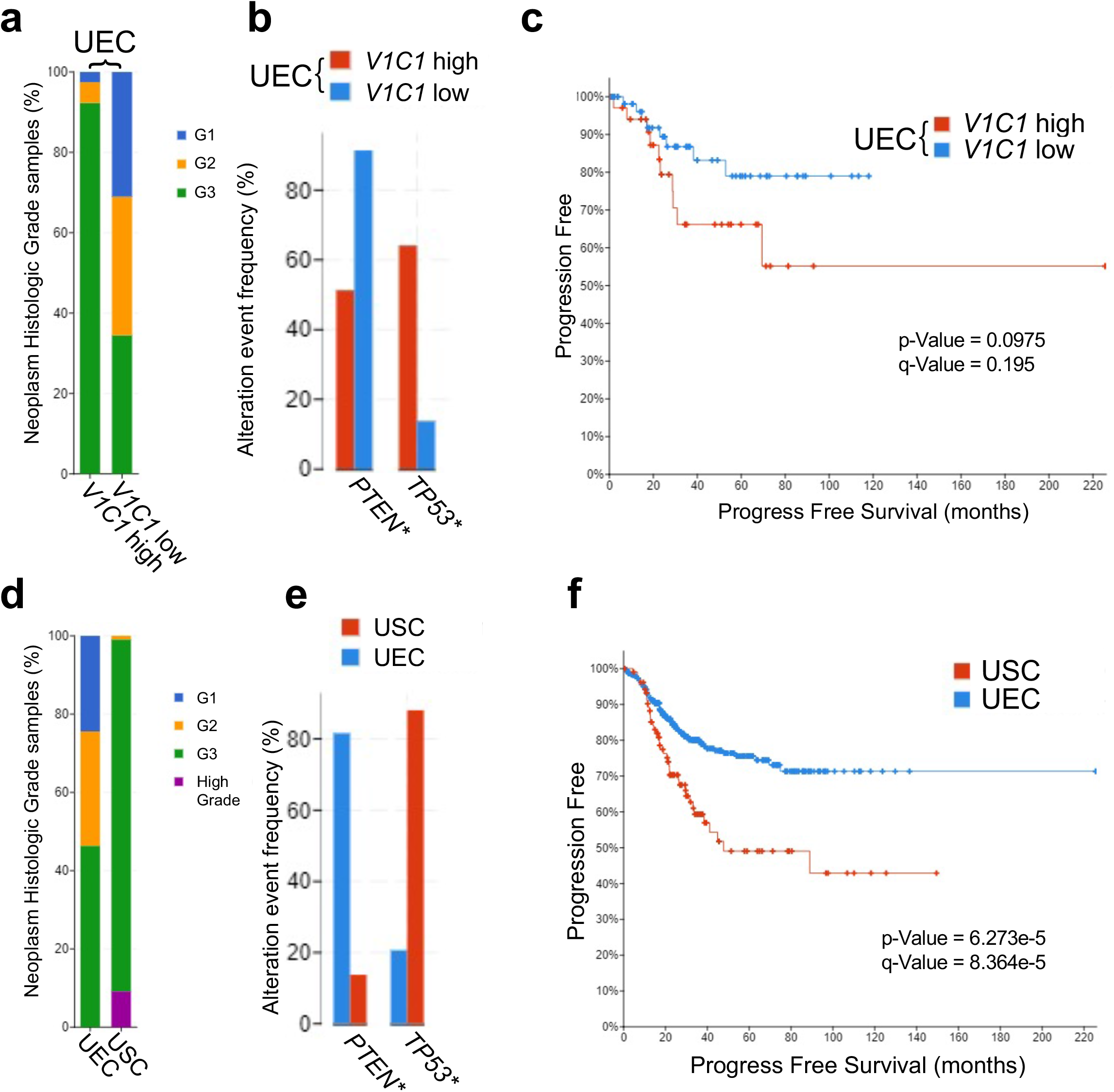
Association between the expression of the V-H^+^-ATPase *C1* isoform subunit, grading, and survival in endometrial carcinoma. Analysis of *ATP6V1C1* high and low expression groups in UEC (**a,b,c**); comparison between UEC and USC (**d, e, f**). Tumor differentiation grade (**a** and **d**); most frequently altered genes in the *ATP6V1C1* high and low expression groups (**b** and **e,** asterisks represent p-value <0.005); Kaplan-Meier curve for progression-free survival in relation to *ATP6V1C1* expression in UEC (**c**) and comparing UEC and USC ((**f)**; log-rank test values are indicated). Abbreviations: PTEN, phosphatase and tensin homologue; TP53, tumor protein p53; UEC, uterine endometrioid carcinoma; USC, uterine serous carcinoma.

Endometrial carcinomas also differ on a molecular basis, i.e., PTEN and PIK3CA which are often mutated in UEC [14], are scarcely altered in USC. On the other hand, mutations of p53 are often observed in USC, but are barely found in UEC (e.g., [1]). Clustering analysis revealed that among the genes clustered with *ATP6V1C1* low expression in UEC, the most frequently mutated gene was PTEN (Fig. 1b), characteristic of tumors with a better prognosis, while mutated TP53, associated with a poor prognosis [1], clustered with C1 high expression group (Fig. 1b; p=1.95e-7), which in turn, exhibit a tumor grade pattern resembling that of USC (Fig.1e). Furthermore, high *ATP6V1C1* expression in UEC could also be associated with molecular malignancy signatures, shared by the USC histological subtype, correlated with highest mortality rates (Fig. 1 a,d). Kaplan-Meier curve of *ATP6V1C1* expression in relation to progression-free survival indicated significant difference between low and high groups, with high *ATP6V1C1* UEC similar to USC profile (Fig. 1 c,f).

Previously, TCGA analyses on the expression of V-ATPase genes in both subtypes of esophageal cancer, squamous cell carcinoma and adenocarcinoma, demonstrated that the expression of C1 isoform was higher in relation to C2a,b isoforms (data validated by qRT-PCR), and that this disproportion between C1 and C2a,b mRNA expressions levels was able to differentiate neoplasia from normal tissues [8]. The present data taken together with our previous multicancer analyses [8], provide compelling evidence for the differential expression of the H^+^ pump subunit C isoforms (especially the dominance of C1) as promising cancer biomarkers. These findings also contributed to the understanding of the role of the V-ATPase complex in tumorigenesis and how this pump can potentiate multiple signaling pathways in coordination with key oncogenes and tumor suppressor genes in UEC [15].

In conclusion, this work opens a new avenue for clinical-pathological studies, which will determine whether this reported molecular phenotyping can be clinically validated and whether expression differences in C1 subunit and other V-ATPase related molecular signatures could be exploited for pathological diagnosis, prognosis and/or targeted therapy, as well as to further understanding of the acid reprogramming metabolism of EC and other related cancers.

## Acknowledgments

This work was supported by the CNPq (Conselho Nacional de Desenvolvimento Científico e Tecnológico), FAPERJ (Fundação de Amparo à Pesquisa do Estado do Rio de Janeiro) and Universidade Estadual do Norte Fluminense Darcy Ribeiro (UENF). This study was financed in part by the Coordenação de Aperfeiçoamento de Pessoal de Nível Superior - Brasil (CAPES) - Finance Code 001.

