## Supplemental Figures S1 and S2 for "Molecular Prognosis of Endometrial Adenocarcinoma by Expression Patterns of the V-ATPase C1 Subunit"

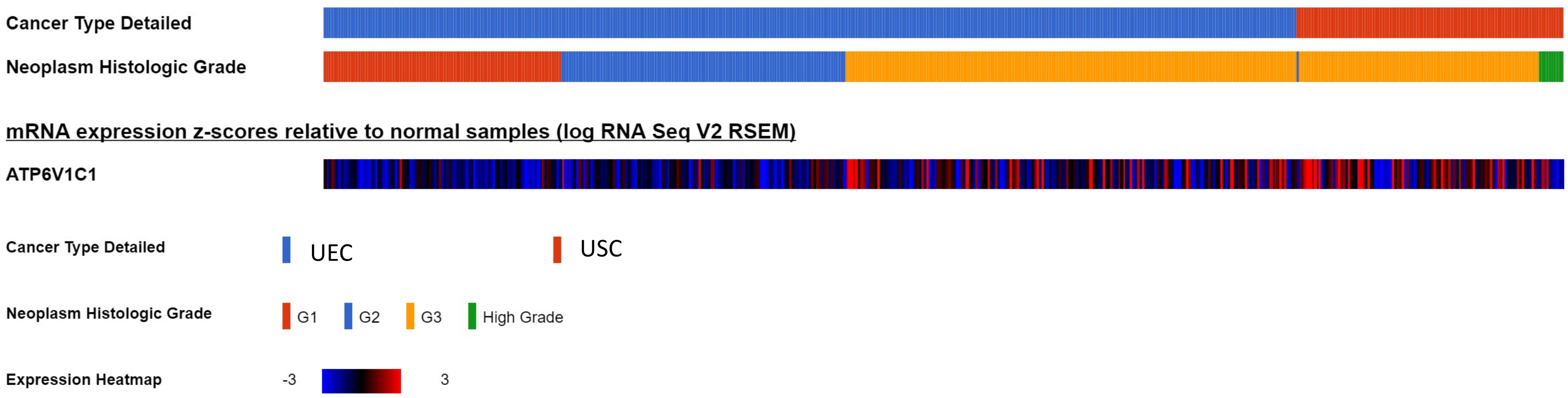


**Supplemental Figure S1.** Expression heatmap of ATP6V1C1 mRNA levels in different histologic grades of uterine endometrioid carcinoma (UEC) and uterine serous carcinoma (USC).

**a**

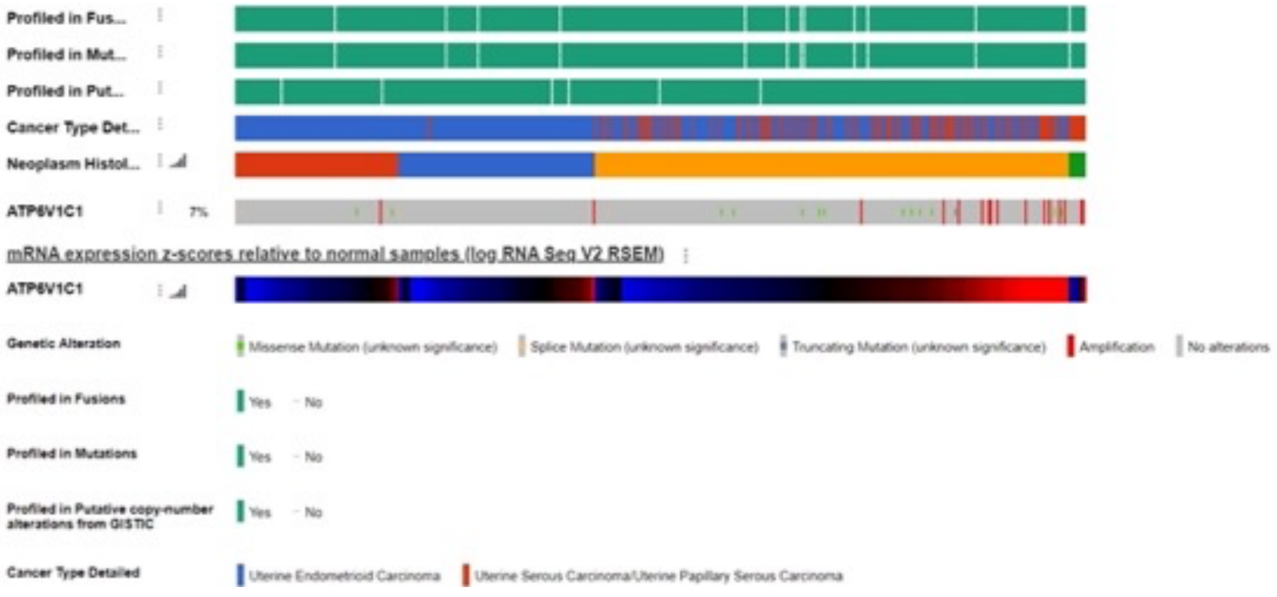


**b**

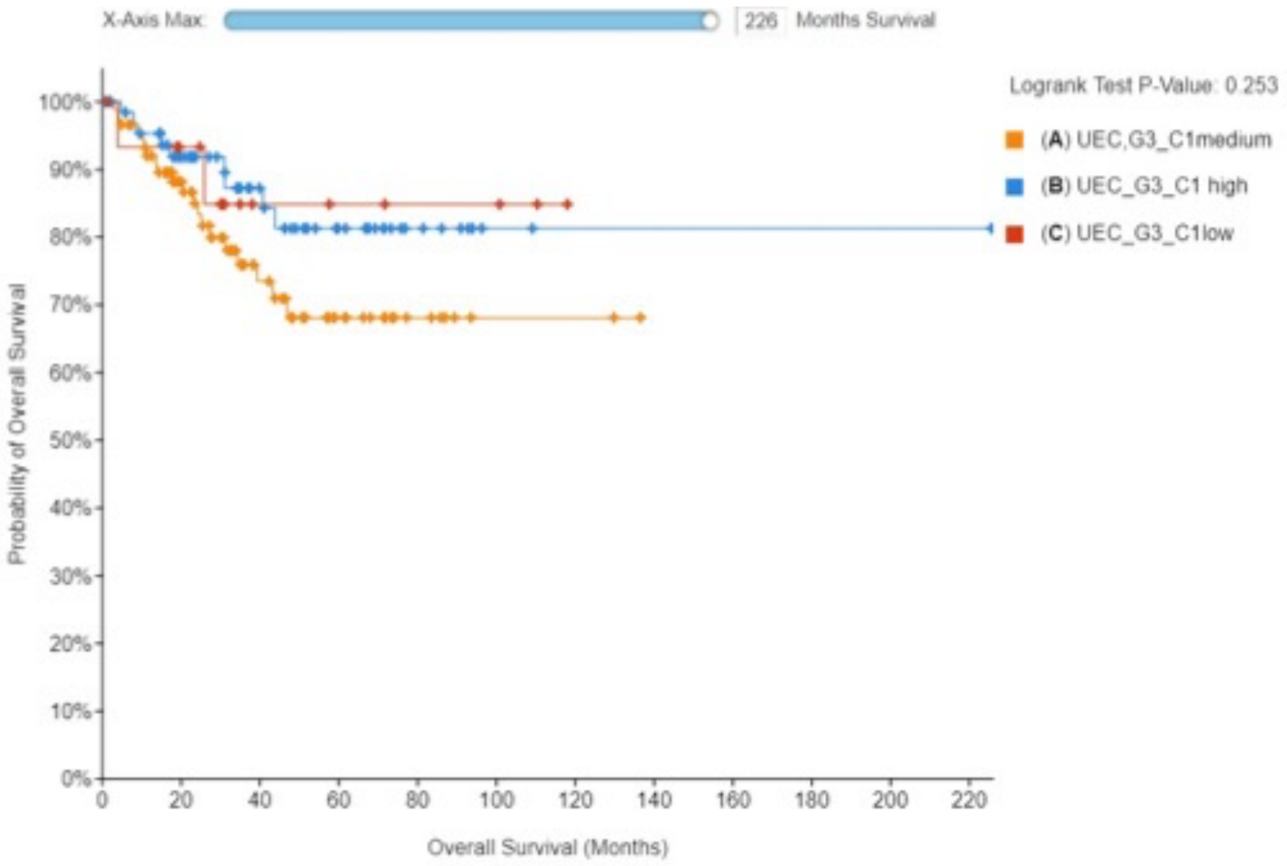


**Supplemental Figure S2.** Relationship between expression and genetic alterations in ATP6V1C1 mRNA and grading in uterine endometrioid carcinoma and uterine serous carcinoma (**a**). Overall survival in G3 uterine endometrial adenocarcinoma in relation to ATP6V1C1 expression levels (**b**).
